# Transgenic pyrimethamine-resistant *P. falciparum* reveals transmission blocking potency of P218, a novel antifolate

**DOI:** 10.1101/2020.09.06.284786

**Authors:** Navaporn Posayapisit, Jutharat Pengon, Parichat Prommana, Molnipha Shoram, Yongyuth Yuthavong, Chairat Uthaipibull, Sumalee Kamchonwongpaisan, Natapong Jupatanakul

**Affiliations:** National Center for Genetic Engineering and Biotechnology (BIOTEC), Pathum Thani, Thailand 12120

**Keywords:** antifolate, drug resistance, malaria, dihydrofolate reductase, transmission blocking, CRISPR-Cas9

## Abstract

Antimalarial drug which target more than one life stage of the parasite are valuable tools in the fight against malaria. Previous generation of antifolate drugs are able to inhibit replicative stages of drug-sensitive, but not resistant parasites in humans, and mosquitoes. The lack of reliable gametocyte-producing, antifolate resistant *P. falciparum* hindrance the development of new antifolate compounds against mosquito stages. We used CRISPR-Cas9 technology to develop transgenic gametocyte producing *P. falciparum* with quadruple mutations in *dhfr* gene, using NF54 as a parental strain. The transgenic parasites gained pyrimethamine resistance while maintaining the gametocyte producing activity. In contrast to pyrimethamine that cannot inhibit exflagellation of the quadruple *dhfr* mutant parasite, the novel antifolate P218 showed a good potency for exflagellation inhibition (exflagellation IC_50_ 10.74 ± 4.22 nM). The exflagellation IC_50_ was 5.3 times lower than erythrocytic IC_50_ suggesting that the human to mosquito transmission poses as a strong barrier to prevent P218 resistant parasite among population. This study demonstrates that P218 can be considered as a highly potent tool to prevent the spread of antifolate resistant parasites.

**Graphical Abstract:** 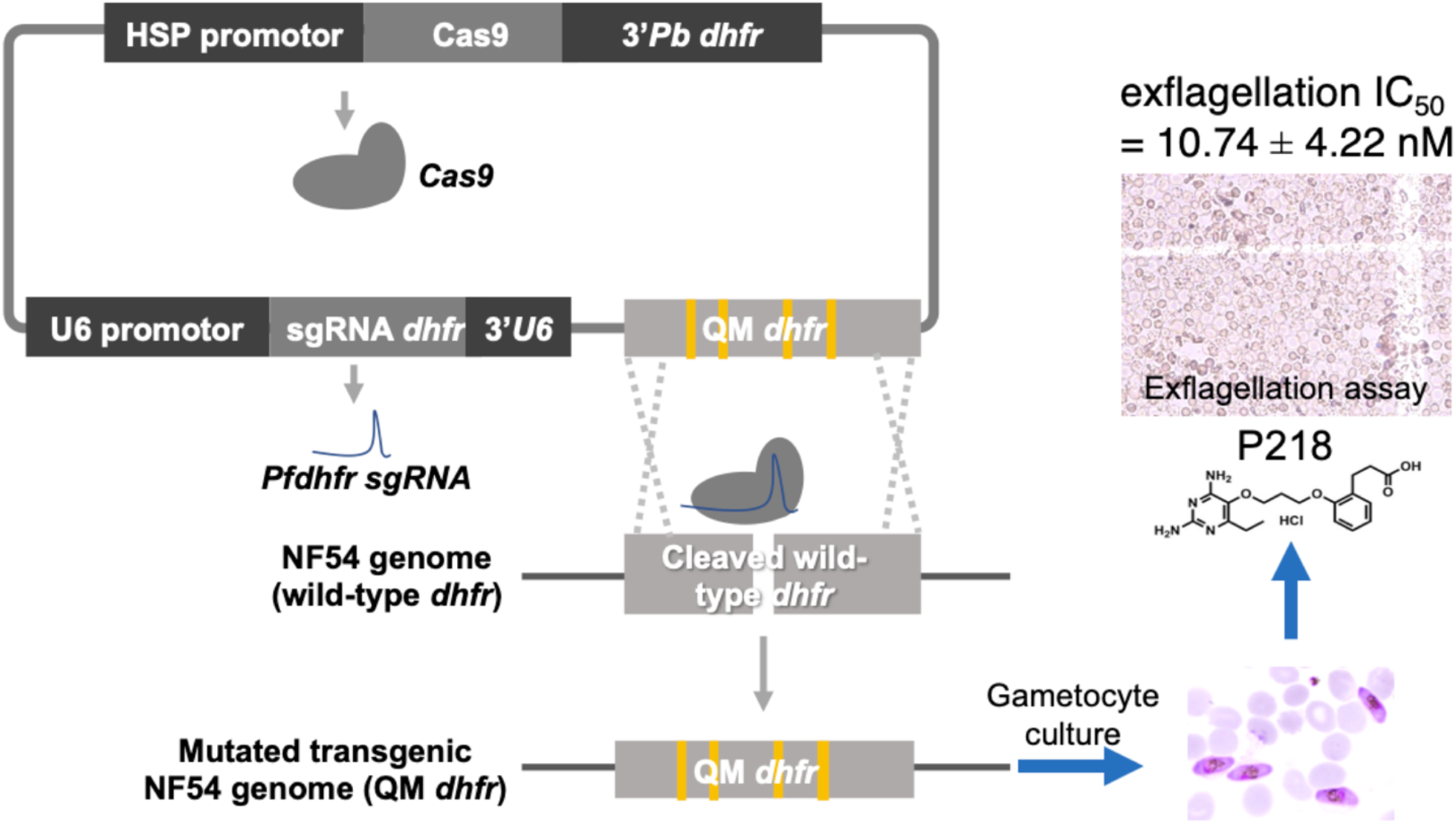

**Research Highlights:**

- Transgenic gametocyte producing pyrimethamine resistant *P. falciparum* was generated.
- P218 asexual stage IC_50_ in NF54-4mut*Pfdhfr* was 56.94 ± 15.69 nM.
- P218 exflagellation IC_50_ in NF54-4mut*Pfdhfr* was 10.74 ± 4.22 nM.
- P218 exflagellation IC_50_ in NF54-4mut*Pfdhfr* is 5.3 times lower than erythrocytic IC_50_.
- P218 is an invaluable tool for malaria treatment and transmission control.

## 1. Introduction

Malaria is a major public health problem that impact many areas of the world causing health, social, and economic impact. The disease is the deadliest mosquito-borne disease causing more than 400,000 deaths from over 200 million cases per year, and most of the death from malaria occur in children under age 5 (World Health Organization, 2019). While effective vaccines are anticipated, antimalarial drugs are still the major means to reduce burden of malaria infection in humans. However, emergence of resistant parasites to the current antimalarials in clinical use has highlighted the importance of antimalarial drug discovery.

During development in the erythrocytic stage, a small percentage of the malaria parasites differentiate into male and female gametocytes, the mosquito-infective stage, which are taken up by mosquitoes during blood meal. Once inside the mosquito gut, changes in environment triggers gametocytes activation (Josling & Llinás, 2015). Male gametocytes undergo exflagellation, which consists of three rounds of DNA replication and differentiation to generate motile parasites called microgametes that can swim through blood meal in the mosquito gut to find and mate with activated female gametocytes (macrogamete). Fertilized diploid zygotes then undergo meiosis and cellular differentiation to generate motile and invasive ookinetes, which then invade the mosquito gut epithelium to form oocysts at around 24 hours after infectious blood meal. The number of parasites that survive these processes in mosquito infection is extremely low and can be considered as a major bottleneck in malaria transmission (Smith et al., 2014). Therefore, the control measures targeting these transmission bottlenecks will be highly effective for malaria control. *Plasmodium* oocysts in the mosquito gut undergo endomitosis to generate thousands of sporozoites over the course of two weeks. Mature sporozoites later emerge from the oocysts then migrate and reside in the mosquito salivary glands before a small proportion of them are injected into human hosts when the infected mosquitoes take a new blood meal. For the goal of malaria elimination, ideal drugs should, in addition to being effective in the pre-erythrocytic and erythrocytic stages, also be able to block the transmission from infected humans to mosquitoes (transmission-blocking) and prevent infection from infectious mosquito bites in human (prophylaxis).

The folate pathway, involving in the one-carbon metabolism, is required for DNA and RNA synthesis as well as other metabolic processes. It is essential for parasite development in both human and insect hosts. Antifolate compounds that target dihydrofolate reductase (DHFR) such as cycloguanil (CG) and pyrimethamine (PYR) have long been used in combination with sulfa drugs for treatment and prophylaxis against malaria since 1940s-1950s (A Gregson, 2005). However, the malaria parasites soon developed resistance to these drugs in 1960s and it was shown that the mutation in DHFR gene is a major contributing factor to the parasite’s resistance to DHFR inhibitor (Aric Gregson & Plowe, 2005; Müller & Hyde, 2013; Yuthavong et al., 2006). The crystal structures for the wild-type bifunctional DHFR-TS and the highly PYR-resistant quadruple mutant enzyme (QM) from *P. falciparum* (Yuvaniyama et al., 2003) gave the structural basis of the lower binding affinity of PYR to quadruple and other mutants (Cowman et al., 1988; Foote et al., 1990; Peterson et al., 1988, 1990; Sirawaraporn et al., 1997). This information was used to design P218, a 2,4-diaminopyrimidine with flexible side chain of 2’-carboxyethylphenyl group (Yuthavong et al., 2012). Although the effectiveness of P218 in antifolate-resistant malaria has been established, the ability of the compound to inhibit mosquito stages of antifolate resistant *P. falciparum* has been much less studied. One of the obstacles for such study was that the laboratory maintained quadruple *dhfr* mutant parasites such as the V1/S strain cannot develop into gametocytes.

With the advance in *Plasmodium* genetic manipulation, it is possible to generate drug resistant parasite using molecular techniques. Previous study showed that NF54 strain *P. falciparum* could preserve an ability to develop into gametocytes even after half a year of asexual stage culture, while the 3D7 strain quickly lose the gametocyte producing (Delves et al., 2016). Because the generation of transgenic parasite is a long process, the ability of the NF54 strain to preserve gametocyte producing capability is suitable for the generation of transgenic parasite to study extraerythrocytic stages.

In the present study, we developed transgenic gametocyte producing *P. falciparum* with quadruple mutation in *dhfr* gene to use as a model for antifolate drug testing in the mosquito stage. With CRISPR-Cas9 technology, the wild-type *dhfr* gene of gametocyte producing NF54 strain P*. falciparum* was replaced by quadruple *dhfr* mutant gene with amino acid mutations at N51I, C59R, S108N, and I164L,the same as the V1/S strain. The transgenic parasite gained antifolate resistance as confirmed in the asexual blood stage while maintaining the gametocyte producing activity. Finally, the activity of P218 in transmission blocking was determined by male gametocyte exflagellation assay.

## 2. Materials and Methods

### 2.1 Ethic statement

Human serum and erythrocytes from donors used for *P. falciparum* culture were obtained under the regulation of the Ethics Committee for Human Research, National Science and Technology Development Agency (NSTDA), following an approved protocol (document number 0021/2560).

### 2.2 P. falciparum strains and asexual stage culture

*P. falciparum* strain NF54 (Patient Line E) MRA-1000 was used as a parental strain for transgenic parasite construction. The parasites comprise a gametocyte-producing strain with wild-type *dhfr* gene.

Asexual blood stage of *P. falciparum* was cultured *in vitro* in human O+ erythrocytes and RPMI medium (Gibco Cat. no. 11875) supplemented with 5% heat-inactivated human serum, 0.125% Albumax I (Gibco Cat. no. 11020), 5.94 g/L HEPES (Sigma Cat. no. H4034), 2 g/L glucose (Sigma Cat. no. G7021), 5 g/L hypoxanthine (Sigma Cat. no. H9377), and 40 mg/L gentamycin sulfate at 37 °C under atmosphere of 94% N_2_ + 5% CO_2_ + 1% O_2_.

Asexual blood stage synchronization was performed by sorbitol treatment. Briefly *P. falciparum* infected erythrocytes were incubated with 5 volumes of 5% sorbitol then incubated at 37 °C for 15 minutes. After incubation, infected erythrocytes were harvested by centrifugation at 1,200 x *g* for 3 minutes. Pelleted erythrocytes were washed with 5 mL of prewarmed RPMI medium before sub-culturing in fresh erythrocytes and complete RPMI to desired hematocrit and parasitemia.

### 2.3 Construction of transgenic gametocyte-producing P. falciparum with mutations on dhfr gene

The *P. falciparum* NF54 strain (MRA-1000) was used as a parental strain for transgenic parasite production as it is an efficient gametocyte-producing strain (Figure 1). The pCas.SgDHFR.HR.V1S generated in previous study consist of Cas9 expression cassette under a control of heat shock promotor and *Pbdhfr* terminator from pUF1-Cas9 plasmid (Ghorbal et al., 2014), single-guide RNA (sgRNA) specific to *P. falciparum dhfr* (*Pfdhfr*) under a control of *P. falciparum* U6 promotor and terminator from pL7 plasmid (Ghorbal et al., 2014), and homology repair quadruple *dhfr* mutant from V1/S strain with mutated PAM site to prevent Cas9 cleavage in the transgenic parasite. Sequence of the pCas.SgDHFR.HR.V1S was in the supplementary information 1. Synchronized ring stage of NF54 strain parasite was transfected with 100 μg of pCas.SgDHFR.HR.V1S plasmid as previously described (Somsak et al., 2011). The transfected parasite was then selected using 1 μM pyrimethamine from day 2 for 14 days, then cloned by limiting dilution method in 96-well plates. The *dhfr* gene from the cloned parasite was then amplified with PfDT Nhe Forward primer (5’ GATGCTAGCATGATGGAACAAGTCTGCG 3’) PfDT3 Hind Reverse primer (5’GCAAGCTTTTAAGCAGCCATATCCATTG 3’) then sent for sequencing with PfD779-pCasR primer (5’ GTGACACTATAGAATACTCAAGCTTTGACATGTATCTTTGTCATC 3’). The mutations of DHFR gene were confirmed by DNA sequencing. The resulting transgenic parasite was called NF54-4mut*Pfdhfr*. The schematic representation of this plasmid is given in Figure 1.

**Figure 1:**
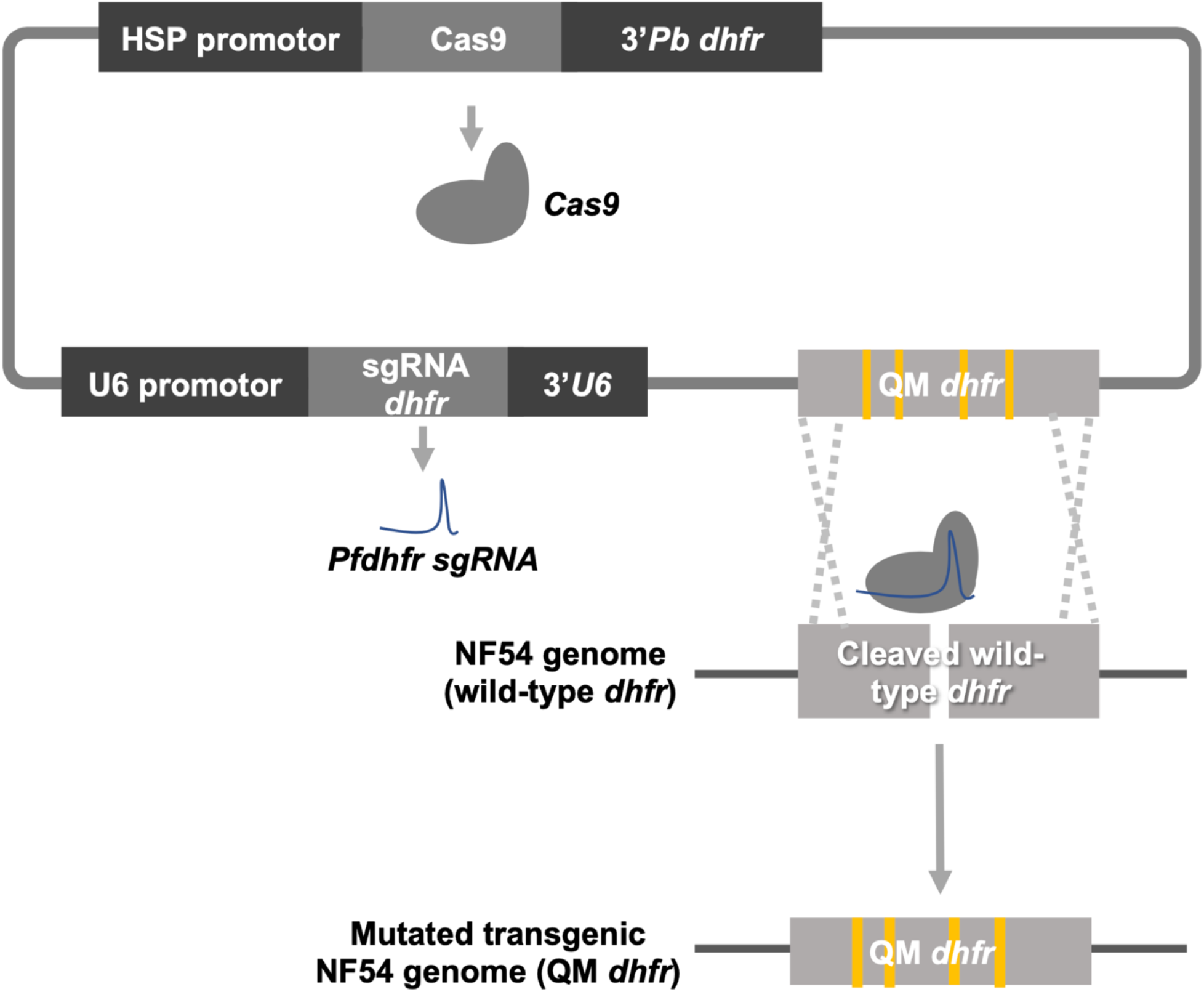
Schematic diagram of the pCas.SgDHFR.HR.V1S plasmid, and the integration of the homology-directed repair cassette into the wild-type *dhfr* locus of the NF54 strain *P. falciparum* to introduce quadruple mutations in the *Pfdhfr* gene.

### 2.4 Assessment of susceptibility of the transgenic NF54-4mutPfdhfr to antimalarial drugs in asexual blood stage

To confirm that the introduction of DHFR mutations to the transgenic NF54-4mut*Pfdhfr* confers antifolate resistance, the transgenic NF54-4mut*Pfdhfr* was tested for its susceptibility to antifolate drugs: pyrimethamine and P218 using Malaria SYBR Green I-base fluorescence (MSF) assay (Johnson et al., 2007). Dihydroartemisinin (DHA) was used as an unrelated drug control. Drug susceptibility of the transgenic parasite was compared with the parental NF54 and V1/S *P. falciparum* parasites. Briefly, 90 μl of 1% ring-stage synchronized parasites at 2% hematocrit were transferred into 96-well flat bottom microtiter plate and treated with 10 μl of serial dilution of each drug prepared in DMSO. The plates were then incubated at 37 °C under atmosphere of 94% N_2_ + 5% CO_2_ + 1% O_2_ for 48 hours followed by an addition of 100 μl of SYBR Green I dye (Invitrogen, Cat. no. S7563) in lysis buffer (20 mM Tris, 5mM EDTA, 0.008% w/v saponin, 0.08% v/v Triton X-100, pH 7.5). Fluorescence signal was measured with a microplate reader (excitation at 435 nm and emission at 535 nm). The SYBR Green I signal from drug-treated parasites were normalized to untreated (DMSO) control parasite in the same experiment.

### 2.5 P. falciparum gametocyte culture

*P. falciparum* gametocytes were cultured following previously published protocol with slight modifications (Delves et al., 2016; Gupta et al., 1985). The culture was started with 1% parasitemia of synchronized ring stage, 4% hematocrit and the medium was changed daily for 16 days. To prevent reinvasion of asexual stage parasite, N-acetyl-glucosamine (NAG, Sigma, Cat. No. A3286) was added to the culture to a final concentration of 50 mM from day 6 to day 11 of the culture. *P. falciparum* gametocyte development was monitored using Giemsa-stained thin blood smear.

### 2.6 Male gamete activation (exflagellation) assay in P. falciparum

The male gamete activation assay was performed following a slight modification of an established protocol (Delves et al., 2013, 2016). In brief, *in vitro* gametocyte culture was set up as described above. On day 13 of the culture, the gametocytes were treated with P218 or other antimalarial compounds at 37 °C for 48 hours. Half of the media supplemented with antimalarial compounds was changed and then incubated for another 24 hours before exflagellation readout. The male gametocyte exflagellation was induced using ookinetes culture medium (RPMI 1640 with 25 mM HEPES, 2 mM glutamine (Sigma, Cat no. G7513), supplemented with 100 μM xanthurenic acid (Sigma, Cat. no. D120804) and observed with hemocytometer under a light-contrast microscope. For each replicate, eight fields of view were recorded with Olympus video camera system (Model DP71) then the total number of exflagellation sites/1,000 red blood cells was calculated then exflagellation inhibition was compared to the DMSO control.

### 2.7 Statistical analyses

Dose response analyses were performed using 4-parameter log-logistic regression model using the *drc* package (Ritz et al., 2015) in the R program (R Core Team, 2020).

## 3. Results

### 3.1 Generation of gametocyte-producing P. falciparum with mutations on DHFR gene

The pCas.SgDHFR.HR.V1S plasmid was successfully transfected into the gametocyte producing NF54 strain. The sequencing results confirmed that the nucleotide at the amino acid position 51, 59, 108, and 164 of the NF54-4mut*Pfdhfr* parasite was successfully mutated (Figure 2).

**Figure 2.**
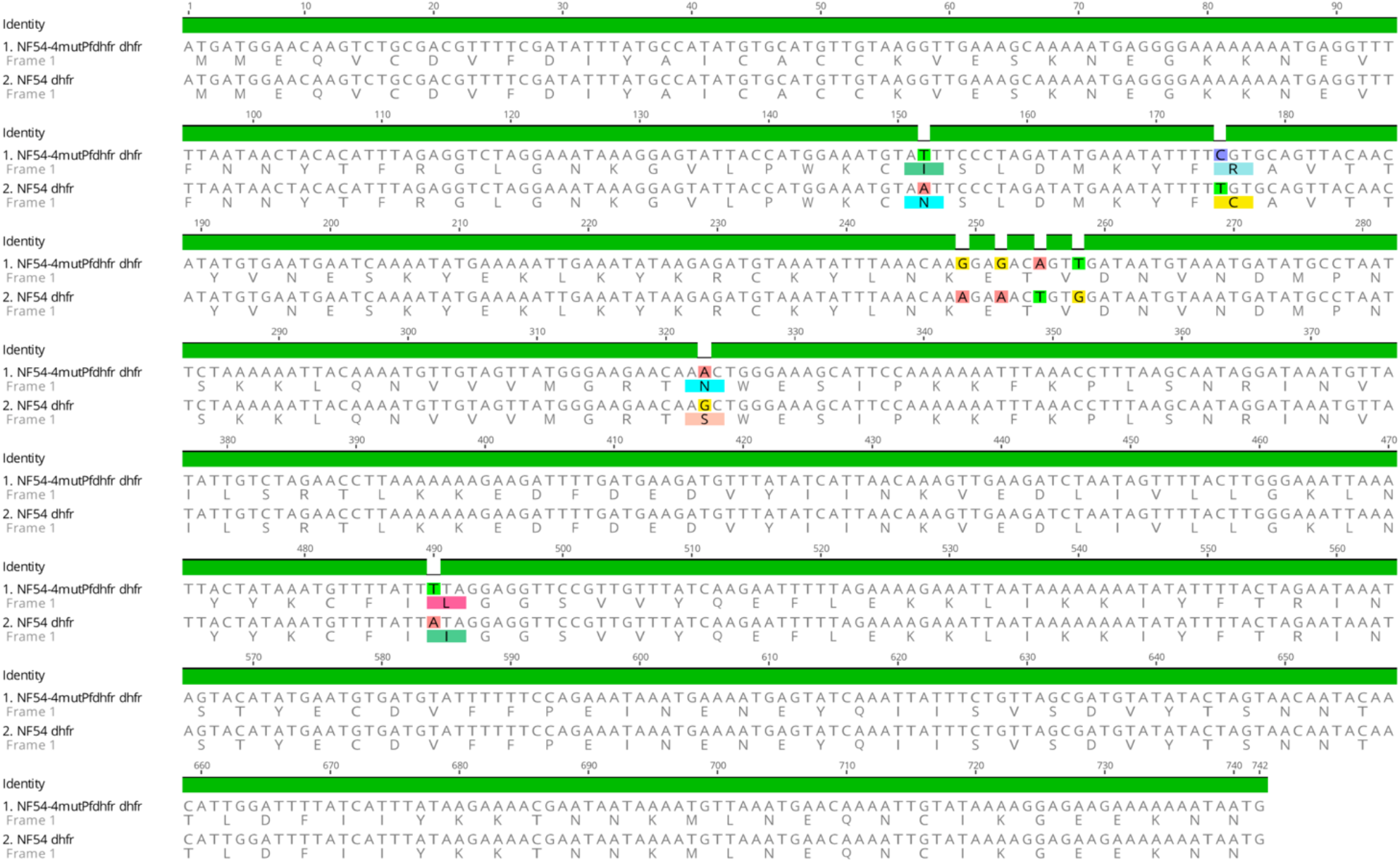
Multiple sequence alignment between the NF54 parental line and the transgenic NF54-4mut*Pfdhfr* parasites. Introduced nucleotide mutations resulted in amino acid mutations at N51I, C59R, S108N, and I164L. Nucleotide 249-258 are synonymous mutations to mutate PAM site for *dhfr* sgRNA.

Because the main goal of generating the transgenic NF54-4mut*Pfdhfr* was to obtain the gametocyte producing parasites with quadruple mutations on the *dhfr* gene, the parasite was then checked for its ability to develop into gametocyte. The NF54-4mut*Pfdhfr* gametocyte was cultured alongside with the NF54 parental and V1/S strain. The result showed that the Nf54-4mut*Pfdhfr* maintain the gametocyte-producing phenotype of the NF54 parental strain with the gametocytemia yield of ~4-10%, which is comparable to the wild-type NF54 strain (Figure 3A). Male gametocytes of the NF54-4mut*Pfdhfr* was highly active and could be further used for exflagellation assay to screen for transmission blocking antifolate compounds (Figure 3B).

**Figure 3.**
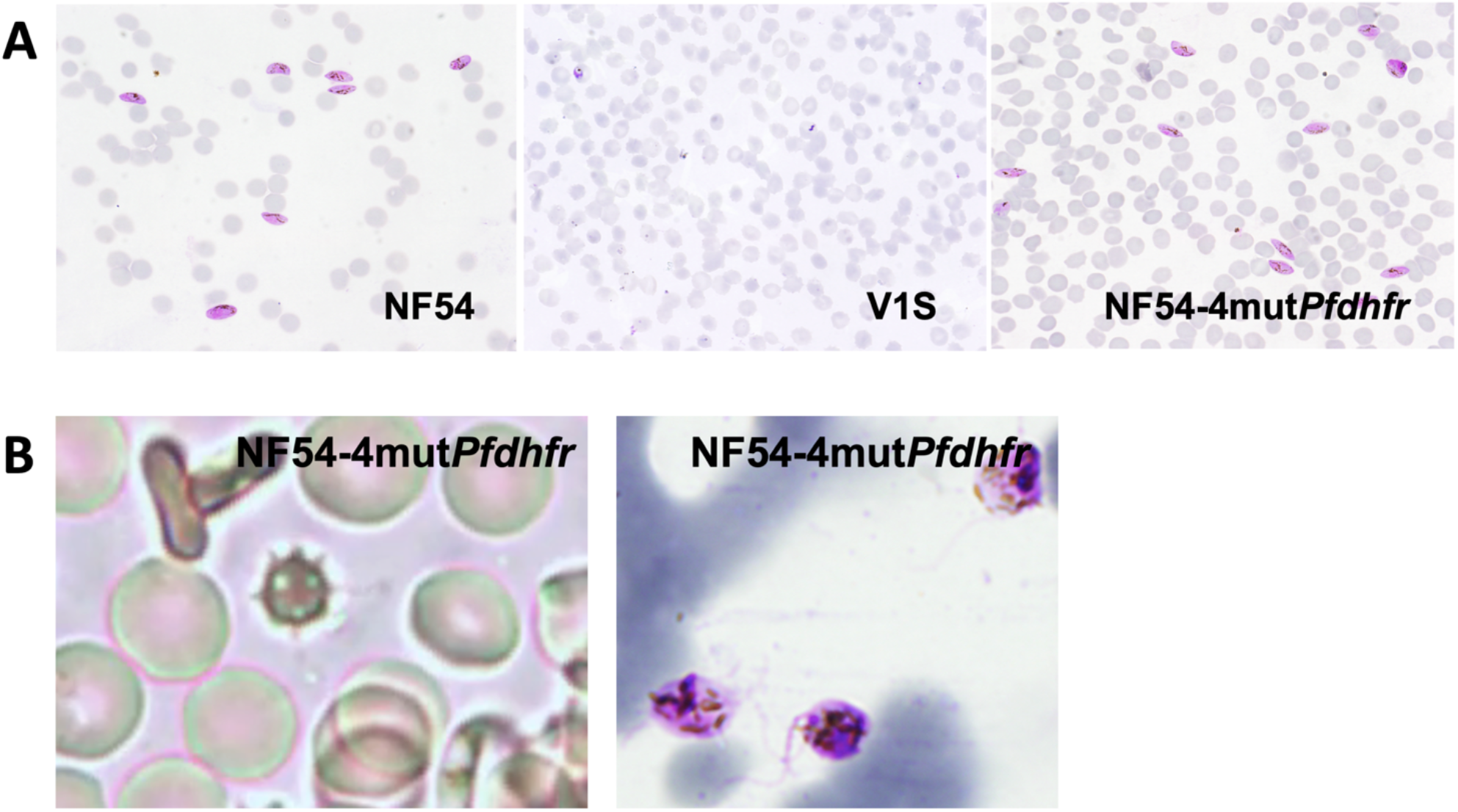
Transgenic NF54-4mutPfdhfr maintains ability to develop into active gametocytes. A) Giemsa stain of thin blood smear of gametocyte culture. B) bright field (left) and Giemsa stained (right) images of activated NF54-4mut*Pfdhfr* male gametocytes.

### 3.2 Asexual erythrocytic stage of the transgenic NF54-4mutPfdhfr has similar antimalarial drug sensitivity compared to the quadruple mutant V1/S strain

After the transgenic parasite was validated and cloned, we then compared its asexual stage antimalarial drug susceptibility to those of the NF54 and V1/S strain (Figure 4, Table 1). While the wild-type NF54 strain is susceptible to PYR with IC_50_ at 126.56 nM, the quadruple mutant V1/S strain had PYR IC_50_ at 65.57 μM or 518-fold higher than that of the wild-type NF54. The replacement of wild-type *dhfr* with quadruple mutant *dhfr* in the transgenic NF54-4mut*Pfdhfr* increased PYR IC_50_ to 87.59 μM or 692-fold increase.

**Table 1.**
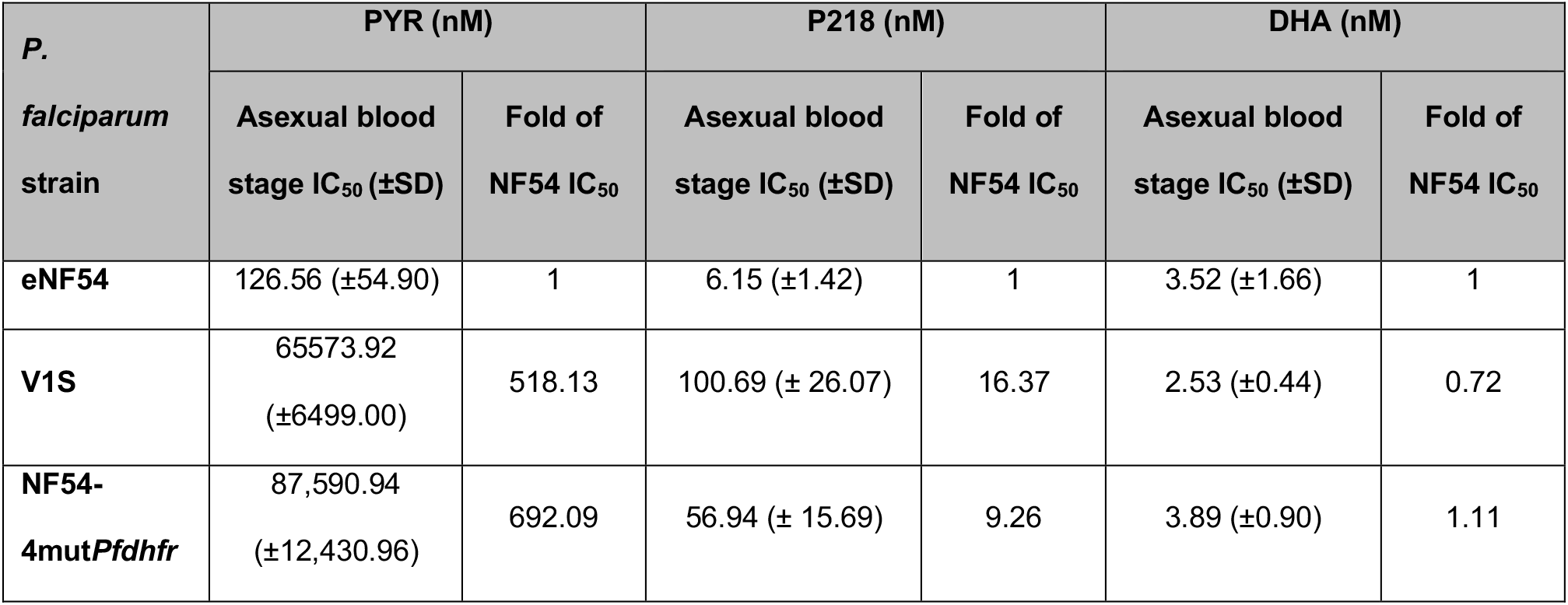
Asexual blood stage IC_50_ of PYR, P218, and DHA in the NF54, NF54-4mut*Pfdhfr*, and V1/S parasites

**Figure 4.**
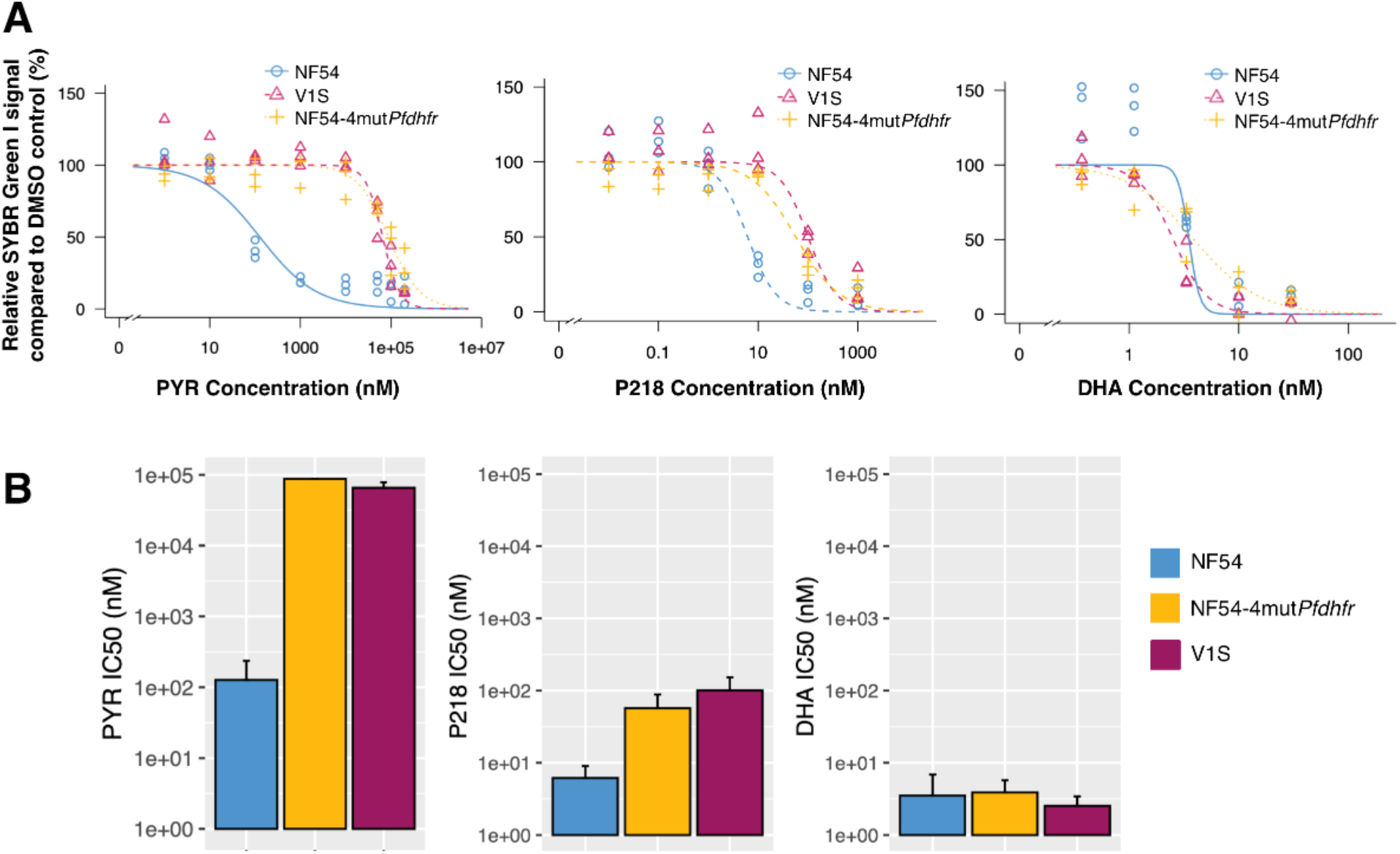
Antimalarial activity of PYR, P218, and DHA in asexual stage of NF54, *NF54-4mutPfdhfr*, and V1/S strains *P. falciparum*. A) Dose response curves of PYR, P218, and DHA. B) Bar charts representing asexual stage IC_50_ values of of PYR, P218, and DHA in NF54, *NF54-4mutPfdhfr*, and V1/S strains *P. falciparum* error bars represent 95% confidence intervals (±2SD) of the IC_50_ values.

Similar to a previous study (Yuthavong et al., 2012), our result demonstrated that P218 has better asexual stage antimalarial activity against both wild-type and quadruple mutant parasites compared to PYR. IC_50_ of P218 in NF54, V1/S, and NF54-4mut*Pfdhfr* were 6.15, 100.69, and 56.94 nM, respectively (Figure 4, Table 1). The increase in IC_50_ correspond to 16- and 9-fold increase for the V1/S, and NF54-4mut*Pfdhfr* strains. These increases are far lower than those for pyrimethamine, with ratios of 32 for V1/S and 75 for NF54-4mut*Pfdhfr*, validating P218 efficacy against pyrimethamine-resistant parasites.

When treated with DHA, an antimalarial compound in different class, the wild-type and transgenic parasites had similar IC_50_ (Figure 4, Table 1). These results demonstrated that the genetic modification specifically increased antifolate resistance.

### 3.3 P218 effectively inhibits male gametocyte exflagellation of the transgenic antifolate resistant parasite

The exflagellation of transgenic and original NF54 strain *P. falciparum* was tested with various antimalarial compounds including PYR, P218, DHA, and methylene blue (MB, an antimalarial compound with confirmed exflagellation inhibition activity) (Figure 5). The exflagellation inhibition of DHA and MB were similar between the NF54 and NF54-4mut*Pfdhfr* parasites suggesting that the replacement of the wild-type *dhfr* to the quadruple mutant *dhfr* only had impact on the parasite’s susceptibility to antifolate compounds (Figure 5, Table 2).

**Table 2.** Exflagellation IC_50_ of PYR, P218, and DHA in the NF54 and NF54-4mut*Pfdhfr* parasites

| *P. falciparum* strain | PYR (nM) |  | P218 (nM) |  | DHA (nM) |  | MB (nM) |  |
| --- | --- | --- | --- | --- | --- | --- | --- | --- |
|  | Exflagellation IC<sub>50</sub> (±SD) | Fold of NF54 IC<sub>50</sub> | Exflagellation IC<sub>50</sub> (±SD) | Fold of NF54 IC<sub>50</sub> | Exflagellation IC<sub>50</sub> (±SD) | Fold of NF54 IC<sub>50</sub> | Exflagellation IC<sub>50</sub> (±SD) | Fold of NF54 IC<sub>50</sub> |
| NF54 | 27.55 (±13.24) | 1 | 4.06 (±1.36) | 1 | 16.39 (±4.43) | 1 | 49.60 (±11.56) | 1 |
| NF54-4mut*Pfdhfr* | 99,885 (±120,920) | 3625.59 | 10.74 (± 4.22) | 2.65 | 20.56 (±8.84) | 1.25 | 32.42 (±17.74) | 0.65 |

**Figure 5.**
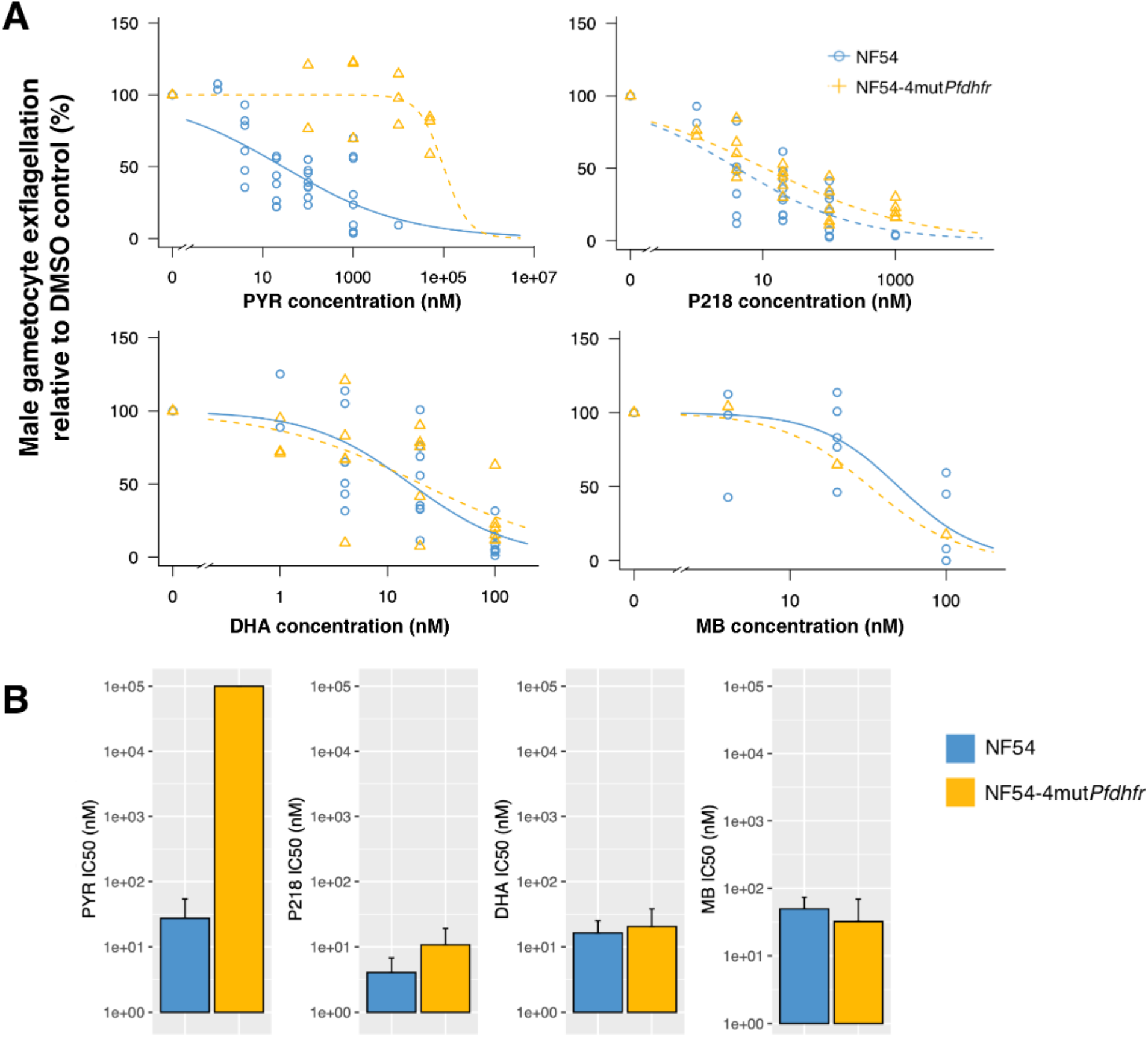
Exflagellation inhibition of PYR, P218, and DHA in NF54 and NF54-4mut*Pfdhfr* strains *P. falciparum*. A) Dose response curve of PYR, P218, and DHA. B) Bar chart representing asexual stage IC_50_ values of PYR, P218, and DHA in NF54 and NF54-4mut*Pfdhfr*. Error bars represent 95% confidence intervals (±2SD) of the IC_50_ values.

As expected, the quadruple *dhfr* mutation substantially increased parasite’s susceptibility to previous generation of antifolate compound such as PYR. PYR exflagellation IC_50_ of the NF54-4mut*Pfdhfr* parasite was 99.89 μM, a 3,626 fold-increase from 27.55 nM in the wild-type *dhfr* NF54 parasite (Figure 5, Table 2). In contrast, exflagellation IC_50_ of P218 in the transgenic quadruple *dhfr* mutant, was 10.74 nM, a 3-fold increase from 4.06 nM in the wild-type (Figure 5, Table 2). Interestingly, the exflagellation IC_50_ of the antifolate compounds were lower than asexual stage IC_50_ except for the PYR IC_50_ in NF54-4mut*Pfdhfr*. This suggests that male gametocyte exflagellation might be more susceptible to antifolate than the asexual blood stage.

## 4. Discussion

In addition to vector control, antimalarial drugs with activity against multiple stages of the parasite life cycle form another key component to achieve the goal of malaria elimination and eradication. Our research group developed novel, rationally-designed antifolate compounds that can inhibit asexual stage of both wild-type and quadruple *dhfr* mutant *P. falciparum* at nanomolar level, with P218 as a lead compound (Yuthavong et al., 2012). However, the activity of P218 against mosquito stage of antifolate resistant parasite, especially for the quadruple mutant parasite, has up to now only been implied without experimental evidence. The laboratory strain of quadruple mutant parasite (V1/S strain) cannot develop into gametocyte stage, thus limiting the transmission blocking activity screening pipeline against quadruple *dhfr* mutant parasite.

To develop a model to determine transmission blocking activity of P218 and other compounds on quadruple *dhfr* mutant parasite, we successfully generated transgenic NF54 strain of *P. falciparum* harboring the *dhfr* gene with quadruple mutations identical to the V1/S strain. The resulting transgenic parasite showed an increased resistance to antifolate drugs in asexual blood stage at a similar level to that of the V1/S parasite while preserving the ability to develop into gametocytes, which can be used for male gametocyte exflagellation assay. The ability of NF54 parasite to maintain gametocyte producing property even after a long transfection and cloning process (up to 2-3 months from transfection until the cloned parasites were obtained) was evident in our study. This demonstrates the possibility to use this approach with other drug targets to study the effect of genetic variations in drug susceptibility of extraerythrocytic stages in the future.

In this study male gametocyte exflagellation was used as a proxy to determine transmission blocking efficiency of P218 because 1) antifolate compounds only target male but not female gametocyte activation (Delves et al., 2013), 2) the exflagellation assay was performed in multi well format *in vitro* thus suitable for testing multiple concentrations for dose response analysis. Our results showed that the exflagellation of male gametocytes are more sensitive to antifolates when compared to the asexual blood stage, which is in concordance with the previously published data of PYR treatment, which reported asexual stage IC_50_ at 17 nM and exflagellation IC_50_ at 8.7 nM (Delves et al., 2013).

Transmission from human to mosquito is a major bottleneck for *P. falciparum* life cycle (Smith et al., 2014). Previous experimental infection of *Anopheles gambiae* mosquitoes with blood *from P. falciparum-infected* patients carrying high gametocytemia (an average of 433.5 gametocytes in each mosquito midgut) resulted in an average of 5.5 ookinete (91.9% prevalence) on day 1, and two oocyst (37.8 % prevalence) on day 7 (Gouagna et al., 1998). With this low number of ookinete and oocyst able to establish infection in the mosquito midgut, the highly potent P218 should drastically prevent malaria transmission from P218-treated patients to mosquito even at nanomolar plasma concentration. This transmission blocking potential remains even for pyrimethamine-resistant parasites, where the exflagellation of which is still effectively inhibited.

Another important aspect from this study is that P218 will be very effective in preventing geographical expansion of drug resistant parasites. Our previous study suggested that additional mutation on *dhfr* gene will impose strong fitness cost to the parasite because the enzyme will have less catalytic activity to natural substrate (Yuthavong et al., 2012). Additionally, even if the parasite gains higher resistance to P218, the higher sensitivity of male gametocyte compared to the erythrocytic stage (exflagellation IC_50_ was 5.3 times lower than erythrocytic IC_50_) will pose a second stronger barrier to prevent the spread of the P218 resistant parasite from that individual to other humans through mosquito. This is a very important aspect for the malaria elimination effort because the compound can prevent the geographical expansion of antifolate resistant parasites.

In summary, this study used CRISPR-Cas9 technology to generate a transgenic gametocyte producing *P. falciparum* with quadruple *dhfr* mutant replacing the wild-type *dhfr.* The parasite was then used to demonstrate that P218, a novel antifolate compound, is highly active against erythrocytic and mosquito stages of *P. falciparum* with quadruple mutations on *dhfr* gene. The compound is thereby an invaluable tool for malaria treatment and transmission control with the goal of malaria elimination.

## Acknowledgements

This work was financially supported by the NSTDA Core Research grant (P-1850116) to SK, and BIOTEC Research unit director initiative grant to NJ (P-1851424) and CU (P-1551235). *P. falciparum* strain NF54 (Patient Line E) MRA-1000 contributed by Megan G. Dowler was obtained through BEI Resources, NIAID, NIH.

## Authors’ Contributions

**Navaporn Posayapisit:** Methodology, Validation, Formal analysis, Investigation, Writing - Original Draft, Writing - Review & Editing

**Jutharat Pengon:** Methodology, Validation, Formal analysis, Investigation, Writing - Original Draft, Writing - Review & Editing

**Parichat Prommana:** Methodology, Validation, Formal analysis, Investigation, Writing - Review & Editing

**Molnipha Shoram:** Investigation

**Yongyuth Yuthavong:** Resources, Writing - Review & Editing

**Chairat Uthaipibull:** Methodology, Resources, Funding acquisition

**Sumalee Kamchonwongpaisan:** Methodology, Resources, Writing - Review & Editing, Funding acquisition

**Natapong Jupatanakul:** Conceptualization, Methodology, Validation, Formal analysis, Investigation, Resources, Writing - Original Draft, Writing - Review & Editing, Visualization, Supervision, Project administration, Funding acquisition

